# External Cues Improve Visual Working Memory Encoding in the Presence of Salient Distractors in Schizophrenia

**DOI:** 10.1101/2021.05.19.442954

**Authors:** Catherine V. Barnes-Scheufler, Lara Rösler, Michael Schaum, Carmen Schiweck, Benjamin Peters, Jutta S. Mayer, Andreas Reif, Michael Wibral, Robert A. Bittner

## Abstract

**Background:** People with schizophrenia (PSZ) are impaired in attentional prioritization of non-salient but relevant stimuli over salient distractors during visual working memory (VWM) encoding. Conversely, guidance of top-down attention by external predictive cues is intact. Yet, it is unknown whether this preserved ability can help PSZ encode more information in the presence of salient distractors.

**Methods:** We employed a visuospatial change-detection task using four Gabor patches with differing orientations in 66 PSZ and 74 healthy controls (HCS). Two Gabor patches flickered which were designated either as targets or distractors and either a predictive or a non-predictive cue was displayed to manipulate top-down attention, resulting in four conditions.

**Results:** We observed significant effects of group, salience and cue as well as significant interactions of salience by cue, group by salience and group by cue. Across all conditions, PSZ stored significantly less information in VWM than HCS. PSZ stored significantly less non-flickering than flickering information with a non-predictive cue. However, PSZ stored significantly more flickering and non-flickering information with a predictive cue.

**Conclusions:** Our findings indicate that control of attentional selection is impaired in schizophrenia. We demonstrate that additional top-down information significantly improves performance in PSZ. The observed deficit in attentional control suggests a disturbance of GABAergic inhibition in early visual areas. Moreover, our findings are indicative of a mechanism for enhancing attentional control in PSZ, which could be utilized by pro-cognitive interventions. Thus, the current paradigm is suitable to reveal both preserved and compromised cognitive component processes in schizophrenia.

## INTRODUCTION

Visuospatial working memory (VWM) and attention are two intricately and bi-directionally related cognitive domains (Cowan, 2001; Gazzaley & Nobre, 2012; Mayer et al., 2007) essential for elucidating the basis of perturbed information processing in schizophrenia (Carter & Barch, 2007). The role of top-down attention for the encoding of VWM representations is particularly important due to clear behavioral (Hahn et al., 2010; Hartman, Steketee, Silva, Lanning, & McCann, 2003; Mayer & Park, 2012) and neurophysiological (Bittner et al., 2015; Haenschel et al., 2007) evidence for a primary impairment of this VWM component process in people with schizophrenia (PSZ).

Specifically, top-down attention is crucial for selecting goal-relevant information to be stored in VWM, and the efficiency of this mechanism has a considerable influence on VWM capacity (Cowan & Morey, 2006; Vogel, McCollough, & Machizawa, 2005). During VWM encoding, top-down attention-which is driven by stimulus relevance, competes with bottom-up attention-which is driven by stimulus salience (Liesefeld, Liesefeld, Sauseng, Jacob, & Müller, 2020; Lorenc, Mallett, & Lewis-Peacock, 2021). Both the salience and relevance of an object influence its processing in VWM (Constant & Liesefeld, 2021). If multiple items compete for limited attentional resources, selection of relevant and inhibition of irrelevant information depends on the successful execution of two top-down control processes (Desimone & Duncan, 1995). First, the control of selection aids the identification of relevant information which should be selected (Luck & Gold, 2008). Second, the implementation of selection enables the differential processing of relevant and irrelevant information (Luck & Gold, 2008). Neural computations during attentional competition assign a distinct priority to each stimulus, which is the combined representation of its salience and behavioral relevance (Fecteau & Munoz, 2006). To this end, the control of attentional selection is guided by an attentional set, which induces a top-down attentional bias toward the most relevant information based on current goals (Bichot, Rossi, & Desimone, 2005; Corbetta & Shulman, 2002; Nicholas Gaspelin & Steven J Luck, 2018; Wolfe, 1994). According to the signal suppression hypothesis, this mechanism also facilitates the active suppression of automatic attentional capture by visually salient distractors through inhibitory mechanisms (N. Gaspelin & S. J. Luck, 2018; Sawaki & Luck, 2010). Thus, generating a priority map involves the parallelized up- and down-weighting of stimuli, both of which are essential elements of the implementation of selection (Nicholas Gaspelin & Steven J Luck, 2018).

Behavioral VWM experiments using flickering stimuli as part of an encoding array clearly indicate a strong attentional bias in PSZ toward highly salient stimuli, even if they are behaviorally irrelevant (Hahn et al., 2010). This bottom-up attentional bias surpasses the top-down bias induced by the attentional set, leading to a failure to correctly prioritize non-salient but relevant stimuli over salient but irrelevant distractors. Consequently, distractors are encoded into VWM with inadequately high priority. The correlation between the magnitude of the bottom-up attentional bias and reduced VWM capacity (Hahn et al., 2010) underscores its importance for VWM dysfunction. These findings indicate a specific impairment of the implementation of attentional selection in PSZ, when top-down control is required to overcome salient distractors. Importantly, control of attentional selection relying solely on an internally generated attentional set is also disturbed in PSZ (Fuller et al., 2006).

Conversely, patients’ ability to utilize external spatial cues to guide top-down attention during VWM encoding and to prioritize information correctly without the presence of salient distractors is intact (Gold et al., 2006). This is indicative of an ‘island of preserved cognitive function’ (Gold, Hahn, Strauss, & Waltz, 2009), which can provide important clues about the cognitive and neurophysiological mechanisms underlying attentional and VWM encoding dysfunction. However, it remains unknown whether top-down attentional control of PSZ aided by external cues can prevail when directly challenged by highly salient distractors. Elucidating this issue is crucial for models of impaired attentional control and stimulus prioritization during VWM encoding. In this context, impaired inhibition due to disturbances in the GABAergic system – a central pathophysiological mechanism in schizophrenia (Howes & Shatalina, 2022; Lewis, Hashimoto, & Volk, 2005) – is highly pertinent.

There is converging evidence that the suppression of attentional capture by salient stimuli relies on top-down inhibitory mechanisms, which induce local inhibition of such stimuli in early visual areas (Nicholas Gaspelin & Steven J Luck, 2018). This process is primarily mediated by inhibitory GABAergic interneurons (Buschman & Kastner, 2015; Zhang et al., 2014). Notably, post-mortem studies have consistently demonstrated widespread abnormalities in cortical inhibitory GABAergic interneurons in PSZ resulting in reduced inhibition (Lewis, 2014). Neuroimaging, behavioral and computational data also support a central role of reduced GABAergic inhibition for cognitive dysfunction in schizophrenia (Shaw et al., 2020). Therefore, impaired GABAergic inhibition in early visual areas might be a crucial mechanism underlying the insufficient suppression of attentional capture by salient stimuli.

Yet, the successful use of spatial cues by PSZ to overcome this deficit would indicate that enhancing inhibitory control of attentional selection exerted by frontal-parietal areas could sufficiently strengthen local inhibition within early visual areas. Moreover, it would imply that impaired control of selection is the central attentional deficit. The goal of the current study was to test these hypotheses in a behavioral VWM experiment directly contrasting top-down and bottom-up attentional processes during VWM encoding. To this end, we employed a VWM paradigm with an encoding array containing an equal number of salient (flickering) and non-salient (non-flickering) Gabor patches with different orientations. Depending on the specific task condition, either the salient or non-salient Gabor patches would be most relevant and probed preferentially (Nicholas Gaspelin & Steven J Luck, 2018). The encoding array was preceded by either a predictive external cue indicating the location of task-relevant stimuli, or a non-predictive external cue providing no such information.

## METHODS AND MATERIALS

### Participants

We recruited 66 PSZ from psychiatric outpatient facilities in and around Frankfurt am Main, Germany. We established diagnoses of all patients according to DSM-5 criteria based on a clinical interview and careful chart review at a consensus conference. We used the Positive and Negative Syndrome Scale (PANSS) to assess current psychopathology. All patients were on stable antipsychotic medication for at least one month at the time of the study.

74 healthy control subjects (HCS) were recruited by online and printed advertisements. They were screened using the German version of the Structural Clinical Interview SCID-I from the Diagnostic and Statistical Manual, Version IV (Saß, Wittchen, Zaudig, & Houben, 2003). They reported no history of any psychiatric illness and no family history of psychiatric illness in first-degree relatives. All participants reported no history of neurological illness, no drug use (excluding nicotine) within the past six months, normal or corrected-to-normal vision, and no color-blindness.

We matched groups for age, sex, premorbid IQ, years of education, and parental years of education (Table 1). Premorbid verbal intelligence was assessed by the German Mehrfachwahl-Wortschatz-Intelligenz Test (Lehrl, Merz, Burkhard, & Fischer, 2005). The ethics committee of the University Hospital Frankfurt approved all study procedures. Subjects provided written informed consent after receiving a complete description of the study.

**Table 1.** Values are mean (SD), or n. All statistics reported are 2-tailed. Abbreviations: HCS = healthy control subjects, PSZ = people with schizophrenia, PANSS = Positive and Negative Syndrome Scale, FGA = first-generation antipsychotics, SGA = second-generation antipsychotics, df = degrees of freedom.

|  | **HCS (n = 74)** | **PSZ (n = 66)** | **Statistic** | **df** | ***p*** |
| --- | --- | --- | --- | --- | --- |
| **Demographics** |  |  |  |  |  |
| Age, Years | 36.70 (13.02); range 19-61 | 37.59 (10.45); range 20-57 | $t = 0.48$ | 138 | 0.656 |
| Sex, Male/Female | 45 / 29 | 44 / 22 | $X^2 = 0.517$ | 1 | 0.294 |
| Participant Education, Years | 16.26 (2.12) | 15.76 (2.77) | $t = -1.19$ | 138 | 0.238 |
| Parental Education, Years | 14.93 (3.02) | 14.82 (3.31) | $t = -0.21$ | 138 | 0.831 |
| Premorbid IQ | 113.93 (11.16) | 109.89 (13.40) | $t = -1.95$ | 138 | 0.054 |
| Schizophrenia |  | 44 |  |  |  |
| Schizoaffective disorder |  | 22 |  |  |  |
| **Psychopathology** |  |  |  |  |  |
| PANSS Positive |  | 10.13 (3.53) |  |  |  |
| PANSS Negative |  | 11.72 (3.83) |  |  |  |
| PANSS General |  | 22.78 (4.85) |  |  |  |
| PANSS Total |  | 44.47 (9.81) |  |  |  |
| **Medication** |  |  |  |  |  |
| FGA | 0 | 1 |  |  |  |
| SGA | 0 | 58 |  |  |  |
| FGA + SGA | 0 | 5 |  |  |  |
| Lithium | 0 | 3 |  |  |  |
| Valproate | 0 | 2 |  |  |  |
| Antidepressants | 0 | 15 |  |  |  |

### Working Memory Task

A visuospatial change detection task using Gabor patches (Figure 1) was implemented on a personal computer using Presentation software version 14.9 (www.neurobs.com). Stimuli were presented on a grey background (RGB values: 191, 191, 191) in a dimly lit room with a viewing distance of approximately 60 cm.

**Figure 1.**
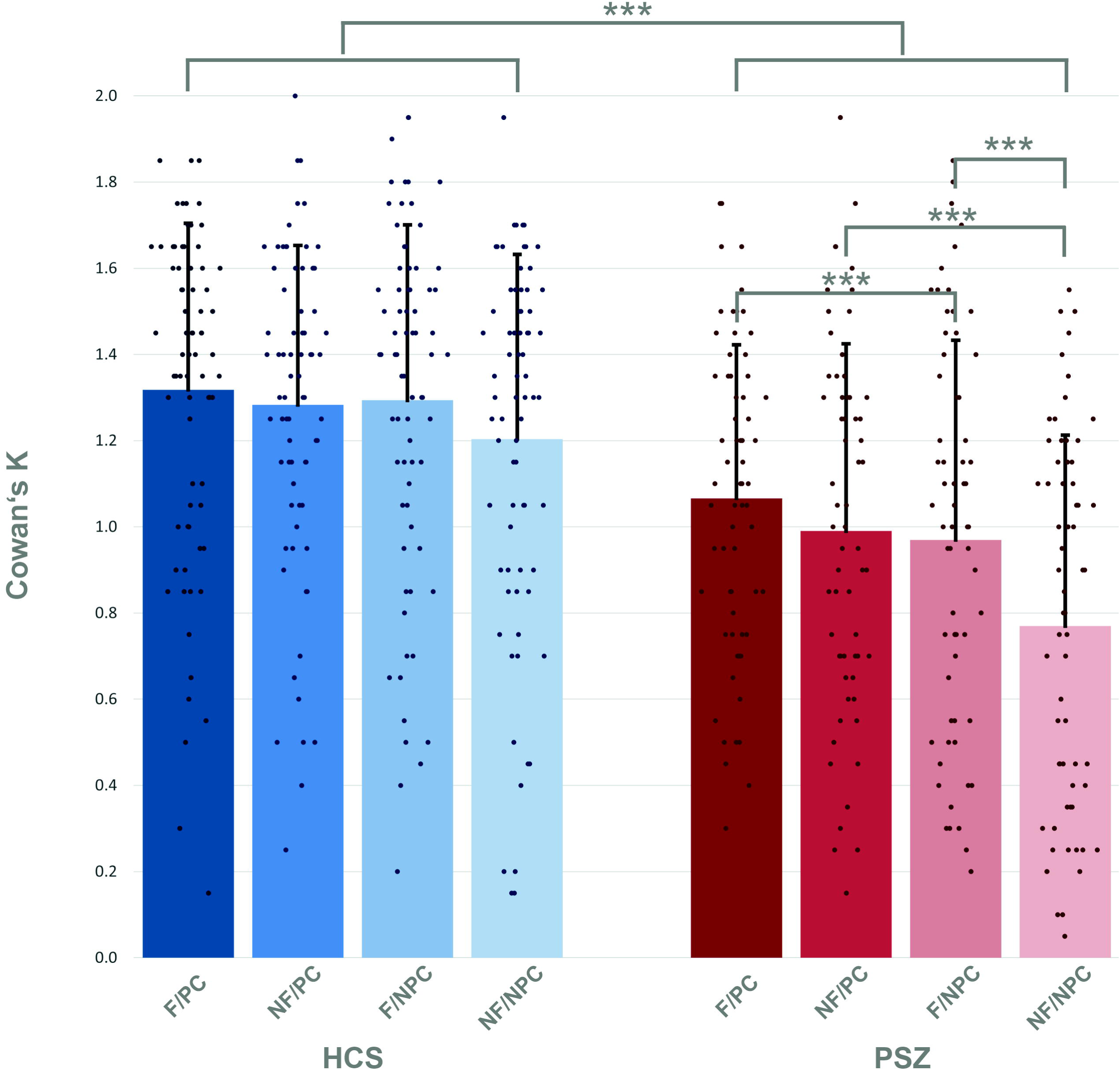
Visual change detection task with four conditions; flickering/predictive cue, flickering/non-predictive cue, non-flickering/predictive cue, non-flickering/non-predictive cue. Flickering is indicated by white dashes around stimuli. The set size of four items was kept constant. In 80% of trials, a designated target stimulus was probed during retrieval (target trials). In 20% of trials, a distractor was probed during retrieval (catch trials). Only target trials are depicted in the figure.

Gabor patches had a visual angle of 0.96° each and were placed on an imaginary circle (visual angle of the diameter: 3.82°). Throughout the experiment a black fixation cross (visual angle: 0.48°) was presented at the center of the screen. Subsequently, either a predictive or a non-predictive external cue was presented by briefly turning the fixation cross white for 300 ms. For predictive cues, the white arms of the fixation cross indicated the future locations of the two most relevant Gabor patches. For non-predictive cues, the entire fixation cross turned white, providing no location information. After a 300 ms preparation interval, the encoding array was displayed for 400 ms consisting of four Gabor patches shown at four fixed locations equally spaced on an imaginary circle around the fixation cross. Two of the four patches flickered at a frequency of 7.5 Hz. To manipulate stimulus relevance, participants were instructed before each block that either the flickering or non-flickering stimuli would be probed preferentially (flickering-bias or non-flickering-bias). Accordingly, in 80% of trials we probed one of the relevant Gabor patches (target trials). In 20% of trials, we probed one of the irrelevant Gabor patches (catch trials). Predictive cues always indicated the future locations of the target Gabor patches irrespective of trial type.

The delay phase lasted for 2000 ms on average, with a jitter of +/- 250 ms. The retrieval array was displayed for 3000 ms consisting of one Gabor patch surrounded by a white frame at the probed location and three blurred out Gabor patches. Probed locations were randomized but counterbalanced. ‘Change’ trials featured a minimum Gabor patch rotation deviation of 45°. Participants had to indicate by button press within 3000 ms, if the orientation of the framed Gabor patch was identical to or different from the Gabor patch shown at the same location in the encoding array. This resulted in four main conditions: flickering-bias/predictive cue, flickering-bias/non-predictive cue, non-flickering-bias/predictive cue, non-flickering-bias/non-predictive cue. Incorporating catch trials in each condition allowed us to assess the efficiency of attentional control operationalized as the difference between the amount of relevant and irrelevant information stored in VWM. A total of 400 trials were presented, 100 for each condition (80 target and 20 catch trials), divided into eight blocks, counterbalanced in order across participants and groups. Each block began with four practice trials to ensure familiarization with the current stimulus relevance and cue type. To obtain an independent estimate of VWM capacity, we also employed a 60-trial canonical visual change detection task (Barnes-Scheufler et al., 2021) (see supplementary materials).

### Analysis of Behavioral Data

The amount of information stored in VWM was quantified using Cowan’s K, where K = (hit rate + correct rejection rate − 1) × memory set size (Cowan, 2001; Rouder, Morey, Morey, & Cowan, 2011). We used a set size of two in accordance with previous studies using comparable task designs (Gold et al., 2006; Hahn et al., 2010). This set size was the most appropriated given that only two stimuli were explicitly designated as targets, leading to a theoretical maximum K of two. Statistical analyses were conducted using SPSS (IBM) Version 22, and R Version 4.3.0 (www.r-project.org). Using a binomial expansion on the 400 trials, we determined that an accuracy of 56% had a cumulative probability of P(X > x) 0.009, i.e., the probability of getting 224 correct responses or more by chance was less than 1%. Accordingly, we excluded participants with an accuracy below 56% (n = 3, all PSZ).

For our main analysis, we fitted a linear mixed model (LMM) estimated using ML and nloptwrap optimizer with R to predict Cowan’s K with group, salience, cue, age, and premorbid IQ (Formula: score ~ Salience * Cue * Group + Age + IQ) and subject as a random effect (formula: ~1 | Subject). We chose this approach because our dependent variable included repeated measures due to the mixed conditions of our study design.

We based this analysis solely on target trials to study the differential processing of the most relevant information. Overall significance estimates of the LMM were obtained post-hoc with an ANOVA function using Satterthwaite’s method. Post-hoc contrasts of the LMM were obtained with the emmeans package (Russell V Lenth, 2023), and were computed using the Kenward-Roger degrees-of-freedom method with pairwise t-tests adjusted with the Tukey method. The t-tests were adjusted for age and premorbid IQ, where age was fixed at 37.1 years and IQ at 112-as determined by the function. Cohen’s d was approximated for the effects in the LMM by transforming t-values.

#### Attentional prioritization and independent VWM capacity estimate

To assess the relationship between the ability to prioritize relevant information and VWM capacity, we correlated the efficiency of attentional prioritization (Cowan’s K for target trials minus Cowan’s K for catch trials) using Spearman correlations in each group across all four conditions with an independent estimate of VWM capacity (Pashler’s K, (Pashler, 1988)) derived from our canonical visual change detection task (Barnes-Scheufler et al., 2021).

#### Correlation between target trials and independent WM capacity estimate

Spearman correlations were conducted with the WM estimate of capacity (Pashler’s K) and target trials (Cowan’s K) in each condition, separately for each group. Fisher z-transformation was used to investigate group differences in correlation strengths.

## RESULTS

### Amount of information stored in VWM

Averaged across all target conditions, HCS encoded 1.27 items, corresponding to an accuracy of 82% (Figure 2, Tables S1 and S2). PSZ encoded 0.95 items, corresponding to an accuracy of 74%. HCS encoded more information into VWM in all four target conditions: flickering-bias/predictive cue (mean = 1.32, SD = 0.39), flickering-bias/non-predictive cue (mean = 1.29, SD = 0.41), non-flickering-bias/predictive cue (mean = 1.28, SD = 0.37), non-flickering-bias/non-predictive cue (mean = 1.20, SD = 0.43) than PSZ: flickering-bias/predictive cue (mean = 1.07, SD = 0.36), flickering-bias/non-predictive cue (mean = 0.97, SD = 0.46), non-flickering-bias/predictive cue (mean = 0.99, SD = 0.44), non-flickering-bias/non-predictive cue (mean = 0.77, SD = 0.44).

**Figure 2.**
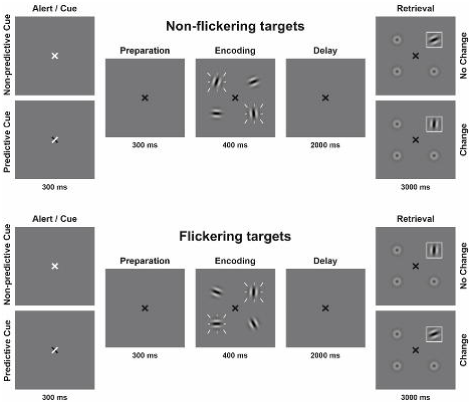
Amount of information stored in VWM in target trials, estimated with Cowan’s K in healthy control subjects = HCS and people with schizophrenia = PSZ. F/PC = flickering/predictive cue, NF/PC = non-flickering/predictive cue, F/NPC = flickering/non-predictive cue, NF/NPC = non-flickering/non-predictive cue. The main group effect of the LMM is expressed as the between group difference, and the within group comparisons are expressed via the post-hoc between group tests. Asterisks indicate significance *p* < 0.001 = ***, *p* < 0.01 = **, *p* < 0.05 = *.Error bars indicate standard deviation.

### LMM effects

We observed significant main effects of group, salience and cue (all *p* < 0.001; Table 2). Additionally, we observed the following significant two-way interactions: group by salience (*p* = 0.037), group by cue (*p* = 0.003), and salience by cue (*p* = 0.013), but no significant three-way interaction between group, salience, and cue (*p* = 0.338). Furthermore, we observed significant effects of the covariates age (*p* < 0.001) and premorbid IQ (*p* = 0.022), but no significant interactions of group by age (*p* = 0.077) or group by premorbid IQ (*p* = 0.354). The model’s total explanatory power was substantial (conditional R^2^ = 0.78), and the part related to the fixed effects alone (marginal R^2^) was 0.25, indicating a better fit when random effects were included (Barton & Barton, 2015).

**Table 2.** Overall significance estimates of the linear mixed model were obtained post-hoc with an ANOVA function using Satterthwaite’s method. Asterisks indicate significance *p* < 0.001 = ***, *p* < 0.01 = **, *p* < 0.05 = *.

|  | **df** | **Sum of Squares** | **Mean Square** | **F** | **p** |
| --- | --- | --- | --- | --- | --- |
| Group | 140 | 1.120 | 1.120 | 24.885 | < 0.001*** |
| Salience | 420 | 1.406 | 1.406 | 31.238 | < 0.001*** |
| Cue | 420 | 1.544 | 1.544 | 34.311 | < 0.001*** |
| Age | 140 | 0.772 | 0.772 | 17.147 | < 0.001*** |
| Premorbid IQ | 140 | 0.241 | 0.241 | 5.360 | 0.022* |
| Group x Salience | 420 | 0.196 | 0.196 | 4.366 | 0.0373* |
| Group x Cue | 420 | 0.394 | 0.394 | 8.763 | 0.003** |
| Salience x Cue | 420 | 0.281 | 0.281 | 6.256 | 0.013* |
| Group x Salience x Cue | 420 | 0.041 | 0.041 | 0.920 | 0.338 |

### Post-hoc tests

To further explore the significant two-way interactions, we conducted post-hoc tests, which revealed a significant reduction of the amount of information stored in VWM in target trials in PSZ compared to HCS in each condition (all *p* < 0.001; Table 3). There were no significant differences between conditions in HCS, however there was a trend level improvement for the amount of stored information in the flickering-bias/non-predictive cue condition compared to the non-flickering-bias/non-predictive cue condition (*p* = 0.050, *d* = 0.2). PSZ stored significantly more information in the flickering-bias/predictive cue condition compared to the flickering-bias/non-predictive cue condition, albeit with a small effect size (*p* = 0.049, *d* = 0.2). Additionally, PSZ stored significantly more information in the non-flickering-bias/predictive cue condition compared to the non-flickering-bias/non-predictive cue condition (*p* < 0.001, *d* = 0.5). Finally, PSZ stored significantly more information in the flickering-bias/non-predictive cue condition compared to the non-flickering-bias/non-predictive cue condition (*p* < 0.001, *d* = 0.5).

**Table 3.**
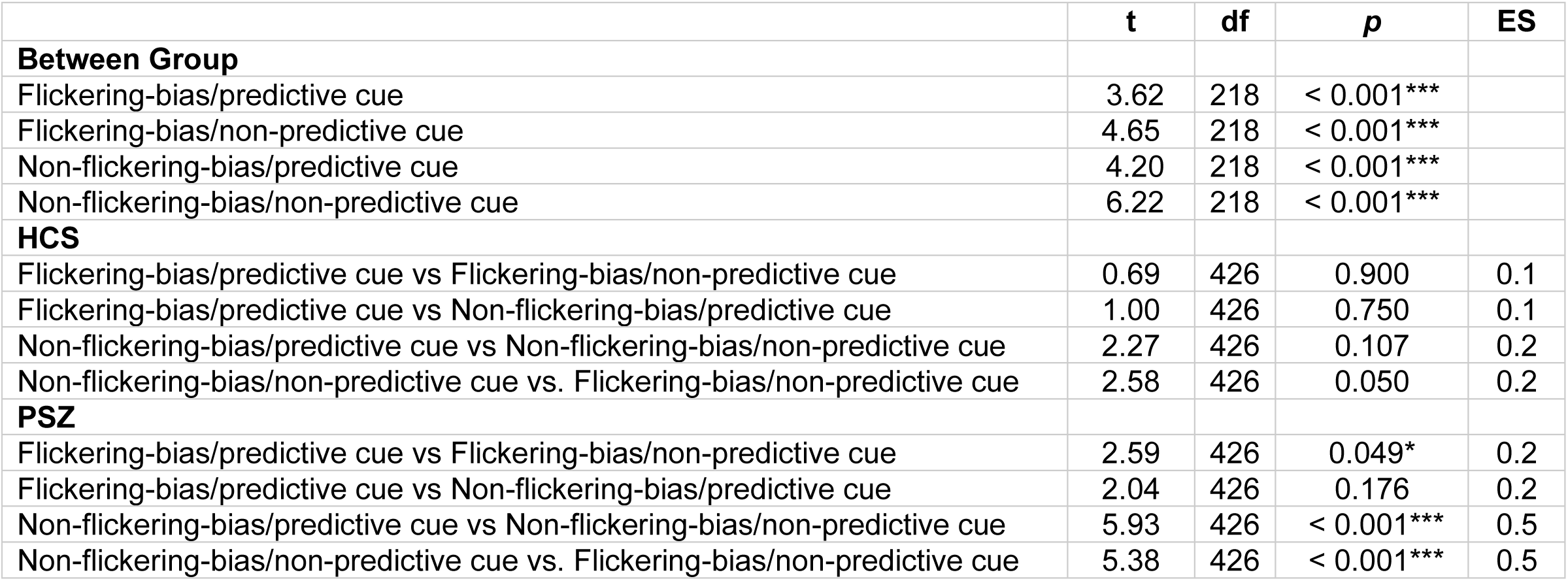
Asterisks indicate significance *p* < 0.001 = ***, *p* < 0.01 = **, *p* < 0.05 = *. Abbreviations: HCS = healthy control subjects, PSZ = people with schizophrenia, df = degrees of freedom, ES = effect size. Comparisons were adjusted for age and premorbid IQ.

|  | **t** | **df** | **p** | **ES** |
| --- | --- | --- | --- | --- |
| **Between Group** |  |  |  |  |
| Flickering-bias/predictive cue | 3.62 | 218 | < 0.001*** |  |
| Flickering-bias/non-predictive cue | 4.65 | 218 | < 0.001*** |  |
| Non-flickering-bias/predictive cue | 4.20 | 218 | < 0.001*** |  |
| Non-flickering-bias/non-predictive cue | 6.22 | 218 | < 0.001*** |  |
| **HCS** |  |  |  |  |
| Flickering-bias/predictive cue vs Flickering-bias/non-predictive cue | 0.69 | 426 | 0.900 | 0.1 |
| Flickering-bias/predictive cue vs Non-flickering-bias/predictive cue | 1.00 | 426 | 0.750 | 0.1 |
| Non-flickering-bias/predictive cue vs Non-flickering-bias/non-predictive cue | 2.27 | 426 | 0.107 | 0.2 |
| Non-flickering-bias/non-predictive cue vs. Flickering-bias/non-predictive cue | 2.58 | 426 | 0.050 | 0.2 |
| **PSZ** |  |  |  |  |
| Flickering-bias/predictive cue vs Flickering-bias/non-predictive cue | 2.59 | 426 | 0.049* | 0.2 |
| Flickering-bias/predictive cue vs Non-flickering-bias/predictive cue | 2.04 | 426 | 0.176 | 0.2 |
| Non-flickering-bias/predictive cue vs Non-flickering-bias/non-predictive cue | 5.93 | 426 | < 0.001*** | 0.5 |
| Non-flickering-bias/non-predictive cue vs. Flickering-bias/non-predictive cue | 5.38 | 426 | < 0.001*** | 0.5 |

#### Attentional prioritization and independent WM capacity estimate

Our independent estimate of WM capacity did not correlate with the efficiency of attentional prioritization across all conditions in HCS (*r_s_* = .162, *p* = 0.167) or in PSZ (*r_s_* = .179, *p* = 0.150), or within any condition of either group (Table 4).

**Table 4.** Results of Spearman correlations (two-tailed) of independent WM capacity estimate (Pashler’s K) and attentional prioritization efficiency (target Cowan’s K – catch Cowan’s K) across all four conditions, and within each condition. Abbreviations: HCS = healthy control subjects, PSZ = people with schizophrenia.

|  | *r<sub>s</sub>* | *p* |
| --- | --- | --- |
| **HCS (*n* = 74 )** |  |  |
| Across all conditions | .162 | 0.167 |
| Flickering-bias/predictive cue | .189 | 0.108 |
| Flickering-bias/non-predictive cue | .155 | 0.187 |
| Non-flickering-bias/predictive cue | .140 | 0.234 |
| Non-flickering-bias/non-predictive cue | .148 | 0.209 |
| **PSZ (*n* = 66)** |  |  |
| Across all conditions | .179 | 0.150 |
| Flickering-bias/predictive cue | .090 | 0.473 |
| Flickering-bias/non-predictive cue | .101 | 0.421 |
| Non-flickering-bias/predictive cue | .101 | 0.421 |
| Non-flickering-bias/non-predictive cue | .212 | 0.087 |

#### Correlation between target trials and independent WM capacity estimate

In HCS, we observed a significant correlation between Cowan’s K in target trials of each of the four conditions and working memory capacity (Pashler’s K) (Table 5). In PSZ, Cowan’s K correlated significantly with working memory capacity (Pashler’s K) only in the two non-predictive cue conditions. However, Fisher z-transformation did not reveal any significant group differences in the strength of these correlations.

**Table 5:**
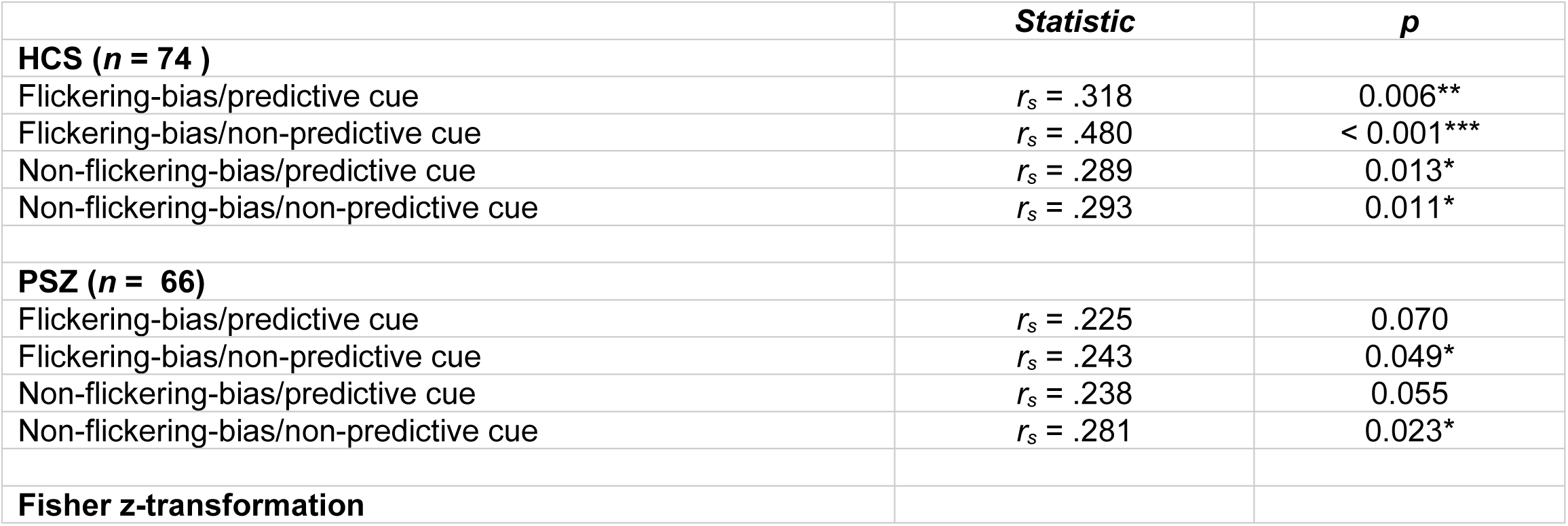
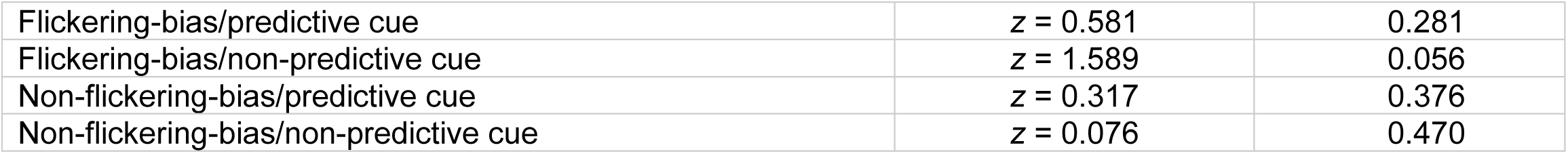
Results of correlations between visual working memory capacity (Pashler’s K) and mean Cowan’s K of each target condition. Asterisks indicate significance *p* < 0.001 = ***, *p* < 0.01 = **, *p* < 0.05 = *. Abbreviations: HCS = healthy control subjects, PSZ = people with schizophrenia. The non-significant z-scores indicate no significant differences between groups of the correlations of VWM capacity and mean target Cowan’s K in each condition.

|  | *Statistic* | *p* |
| --- | --- | --- |
| **HCS (*n* = 74 )** |  |  |
| Flickering-bias/predictive cue | *r<sub>s</sub>* = .318 | 0.006** |
| Flickering-bias/non-predictive cue | *r<sub>s</sub>* = .480 | < 0.001*** |
| Non-flickering-bias/predictive cue | *r<sub>s</sub>* = .289 | 0.013* |
| Non-flickering-bias/non-predictive cue | *r<sub>s</sub>* = .293 | 0.011* |
| **PSZ (*n* = 66)** |  |  |
| Flickering-bias/predictive cue | *r<sub>s</sub>* = .225 | 0.070 |
| Flickering-bias/non-predictive cue | *r<sub>s</sub>* = .243 | 0.049* |
| Non-flickering-bias/predictive cue | *r<sub>s</sub>* = .238 | 0.055 |
| Non-flickering-bias/non-predictive cue | *r<sub>s</sub>* = .281 | 0.023* |
| **Fisher z-transformation** |  |  |
| Flickering-bias/predictive cue | $z = 0.581$ | 0.281 |
| Flickering-bias/non-predictive cue | $z = 1.589$ | 0.056 |
| Non-flickering-bias/predictive cue | $z = 0.317$ | 0.376 |
| Non-flickering-bias/non-predictive cue | $z = 0.076$ | 0.470 |

## DISCUSSION

We studied the competition between bottom-up and top-down attentional processes during VWM encoding in PSZ and HCS to elucidate how disturbed attentional control contributes to VWM dysfunction in schizophrenia. We observed a significant effect of group with a decreased amount of information stored in VWM in PSZ across all conditions. Additionally, we observed a significant effect of salience and a significant interaction of salience by group. With predictive cues, there were no differences in the amount of stored information for salient compared to non-salient targets in either group. However, with non-predictive cues, PSZ stored significantly more information for salient compared to non-salient targets. HCS showed a trend in the same direction. We also observed a significant effect of cue and a significant interaction of cue by group. Only in PSZ did predictive cues increase the amount of information stored in VWM for both salient and non-salient targets compared to non-predictive cues. Notably, this improvement was not sufficient to increase the amount of stored information up to the level of HCS.

Overall, we could replicate the bottom-up attentional bias toward salient distractors during VWM encoding in PSZ (Hahn et al., 2010) and demonstrate that PSZ were able to utilize predictive external cues to overcome this deficit in attentional control. Notably, the use of stationary rotating stimuli as distractors during VWM encoding did not induce a bottom-up attentional bias in PSZ (Erickson et al., 2015). Yet, here stimulus salience might not have been sufficient to elicit automatic attentional capture, especially in light of extensive evidence for motion processing impairments in PSZ (Chen, 2011). Conversely, our flickering visual stimuli should be particularly salient (Merigan & Maunsell, 1993).

However, the central question of our study was concerned with the effects of a predictive cue on attentional prioritization in the presence of salient distractors in PSZ. For the non-flickering-bias conditions, where participants had to suppress highly salient stimuli, a predictive cue significantly increased the amount of stored information only in PSZ. This implies that patients were able to utilize external cues to up-weight the most relevant information. However, there were no group differences in the amount of stored information in catch trials (Figure S2) and thus no evidence for impaired down-weighting of distractors in PSZ. Therefore, contrary to our main hypothesis we did not observe a specific deficit in attentional prioritization in PSZ. Given that PSZ performed significantly worse in the non-flickering/non-predictive cue compared to the flickering/non-predictive cue condition, attentional capture of salient distractors might have still interfered more strongly with the up-weighting of non-salient targets.

Importantly, our task differs from a previous study reporting impaired attentional prioritization in PSZ (Hahn et al., 2010), which directly probed stimulus location. This design likely increased the impact of attentional capture on the encoding of distractor-related information. Conversely, in our study, automatic attentional capture of the locations of distractors might not have sufficiently facilitated the encoding of their orientation. Encoding this additional stimulus feature despite its low relevance would have likely required the voluntary allocation of additional attentional resources. This might have obscured possible deficits in the down-weighting of salient distractors in PSZ.

For PSZ, both high stimulus salience and external cues increased the amount of relevant information stored in VWM. In line with previous studies, these findings indicate that patients’ ability to utilize these specific types of bottom-up and top-down cues to implement attentional selection is largely intact (Gold et al., 2017; Gold et al., 2006). They also confirm that patients’ implementation of selection is impaired, when guided only by the attentional set (Fuller et al., 2006), which appears to contribute to the overall reduction of VWM capacity.

Yet, even when aided by both a predictive cue and high target salience, patients still showed a substantial VWM deficit. Therefore, our findings are compatible with a reduced VWM capacity in schizophrenia independent from specific deficits in attentional control (Barnes-Scheufler et al., 2021; Hahn, Robinson, Leonard, Luck, & Gold, 2018; Leonard et al., 2013). The correlations between target-trial performance and our independent measure of VWM capacity also support this notion, indicating that PSZ might have fewer slots in VWM.

Importantly, impairment in WM capacity appears to be closely connected to global cognitive impairment in PSZ, but can be distinguished from deficits in executive function (Gold et al., 2017; Johnson et al., 2013). Reduced WM capacity in PSZ could be explained by more severely limited attentional weights available for allocation across items. This is compatible with the finding that patients are impaired in their ability to distribute attention broadly (Gray et al., 2014) and show attentional hyperfocusing during visual information processing (Hahn et al., 2022; Luck, Leonard, Hahn, & Gold, 2019). It has been argued that hyperfocusing on internal representations in PSZ might constrain both VWM capacity and attentional resources (Luck, Hahn, Leonard, & Gold, 2019). Therefore, future studies should investigate the relationship between the capability to distribute attention broadly and attentional filtering during VWM encoding.

Moreover, increased vulnerability to attentional capture by salient distractors in PSZ can also be interpreted as a manifestation of attentional hyperfocusing, i.e. hyperfocusing on irrelevant information. This could imply a shared neurophysiological mechanism involving abnormal GABA-mediated inhibition for both phenomena. However, the exact relationship between the impaired implementation of attentional selection in the presence of salient visual distractors and the degree of attentional hyperfocusing in PSZ remains to be investigated. As attentional hyperfocusing appears to constrain the amount of information PSZ can encode into VWM (Luck, Hahn, et al., 2019), it is conceivable that it contributed to the general VWM impairment we observed in PSZ across all task conditions. However, we were not able to make any inferences regarding this interpretation based on our current paradigm.

Our results are well in line with a differential impairment of the two attentional constructs (Luck & Gold, 2008). The control of selection appears to be largely intact in PSZ as the amount of stored information increased significantly when predictive cues were used to guide top-down attention. Conversely, the implementation of selection appears to be impaired in PSZ as less information was stored across all conditions. Post-hoc within group analyses also revealed that our independent estimate of VWM capacity correlated significantly with the amount of stored information in target trials of each condition in HCS, but only in the non-predictive cue conditions in PSZ (Table 5). However, there were no significant group differences regarding the strength of these correlations for any condition as indicated by the Fisher r to z transformation.

Furthermore, disturbances during VWM consolidation, which can constrain performance independent of storage capacity (Xie & Zhang, 2017, 2018), may have also contributed to the overall VWM deficit in patients considering that VWM consolidation appears to be slowed in schizophrenia (Fuller et al., 2009; Fuller, Luck, McMahon, & Gold, 2005; Stablein et al., 2018).

Notably, the analysis of catch trials revealed a significant effect of cue (*p* = 0.004; Table S5). Predictive cues enhanced the suppression of irrelevant information in both groups, suggesting that patients’ ability to use an external spatial cue to down-weight both salient and non-salient distractors is preserved. Additionally, our results are compatible with the interpretation that enhanced inhibitory control during attentional selection increases local inhibition within early visual areas. In accordance with the with the signal suppression hypothesis, increased inhibition could result in a relative enhancement of the cued locations in the attentional priority map by suppressing irrelevant locations.

The balance of excitatory and inhibitory (E/I) activation mediated by interneurons is a central principle underlying cortical computations, and is crucial for the generation of gamma oscillations, which support cognitive processes including attention and working memory (Buzsáki & Wang, 2012; Fries, 2009). Disturbances in the E/I balance in schizophrenia have been attributed to neurobiological and functional abnormalities of cortical GABAergic inhibitory interneurons (Anticevic & Lisman, 2017). Specifically, reduced inhibition of excitatory pyramidal cells can cause widespread disturbances in the E/I balance (Gonzalez-Burgos, Cho, & Lewis, 2015). The bottom-up attentional bias toward salient distractors observed in our paradigm is indicative of reduced inhibition in schizophrenia, implicating GABA-A receptors as potential targets for future pro-cognitive pharmacological interventions (Page & Coutellier, 2018).

Interestingly, there was one significant group difference regarding the efficiency of attentional prioritization (target Cowan’s K minus catch Cowan’s K) for the non-flickering/non-predictive cue condition, with more information being stored by HCS (Table S8). In this condition the efficiency of attentional prioritization was lowest in PSZ, most likely due to hyperfocusing on salient, irrelevant information in the absence of an additional top-down predictive cue.

Notably, we observed a significant correlation of age and the cue effect only in HCS (*r_s_* = 0.240, *p* = 0.040; Table S4), indicating that with increasing age, the difference in the amount of stored information between predictive and non-predictive cue conditions increases. This could provide clues regarding which specific cognitive processes deteriorate in the aging brain. Additionally, in PSZ we observed a significant negative correlation between flickering and non-flickering conditions and premorbid IQ (*r_s_* = −0.254, *p* = 0.039; Table S4). This implies that with increasing IQ, the difference in the amount of stored information between flickering and non-flickering conditions decreases. It appears therefore, that patients with higher IQ were better able to guide attention to the most relevant items irrespective of their salience. This is particularly relevant considering that premorbid IQ predicts functional outcome in PSZ (Leeson, Barnes, Hutton, Ron, & Joyce, 2009).

One limitation of our study is the lack of eye-tracking data to control for group differences in possible deviations of fixation, especially saccades to salient distractors during non-flickering-bias conditions. Both PSZ and HCS performed near chance level during catch trials across all conditions. However, excessive saccades to salient distractors might have still interfered with the encoding of target stimuli.

To summarize, the current paradigm allowed us to evaluate the utilization of external predictive cues to improve WM performance with visually salient distractors in PSZ. We report an overall reduction in the amount of information encoded into VWM across all four conditions in PSZ compared to HCS, with a significant improvement using predictive cues. Our findings are consistent with a limited VWM slot model in PSZ (Barnes-Scheufler et al., 2021), yet also indicate the possibility for a significant improvement in performance with the correct aids. The neurophysiological underpinnings of our findings should be illuminated using functional neuroimaging. The relevance of such a study is also underscored by the inclusion of both attention and working memory as constructs in the cognition domain of the Research Domain Criteria initiative (Insel et al., 2010). Given the presence of deficits in attention and working memory in related neuropsychiatric disorders such as bipolar disorder, our paradigm might therefore be valuable for studying the pathophysiology of impaired information processing across diagnostic categories.

## ACKNOWLEDGEMENTS

The authors are very thankful to Christoph Fehr and Peter Hustedt for their support in recruiting patients, as well as to Deliah Macht, Lisa Goldschmidt, Tobias Lehmann and Caroline Mirkes for assisting with data acquisition, and Danko Nikolic for helpful feedback.

## AUTHORS’ CONTRIBUTIONS

All authors made substantial contributions to the conception or design of the work, or the acquisition, analysis or interpretation of data. Authors LR, MS, BP, JSM, MW, and RAB designed the experiment. Authors CVB-S and LR acquired the data. Authors CVB-S, CS, BP, JSM and RAB analyzed the data. Authors CVB-S and RAB undertook the literature searches and wrote the first draft of the manuscript. All authors contributed to and revised the manuscript. All authors read and approved the final manuscript.

## ETHICAL STANDARDS

The authors assert that all procedures contributing to this work comply with the ethical standards of the relevant national and institutional committees on human experimentation and with the Helsinki Declaration of 1975, as revised in 2008.

