## Supplementary Materials for "External Cues Improve Visual Working Memory Encoding in the Presence of Salient Distractors in Schizophrenia"

### METHODS:

*Demographics:*

Parental Education: Highest value of either parent was used to quantify parental education.

We assessed handedness as a continuous variable using the Edinburgh Handedness Inventory (Oldfield, 1971). We compared handedness scores between groups using a t-test. Spearman Correlations (two-tailed) were used to investigate possible relationships between gender and overall target Cowan's K in both groups.

*Main working memory task:*

Participants were informed about the currently task-relevant Gabor patches (flickering-bias or non-flickering-bias) and of the high likelihood that they would be probed during retrieval. For example, in the flickering-bias predictive cue condition, the instructions read 'In this task, the flickering-bias patterns will be probed preferentially. The positions of these striped patterns will be marked by means of the fixation cross,' and was displayed for 15 seconds.

Accuracy was also calculated as percent correct, and is provided here as a further means of performance evaluation (Table S1). An in-depth breakdown of Cowan's K scores in each condition in each group in both target and catch trials is provided in Table S2.

*Investigation of overall possible influences of psychopathology and medication*

We were also interested in the relationship between the amount of information encoded into working memory and clinical variables. To this end, we performed Spearman bivariate correlations (2-tailed) in PSZ to examine the relationship between total PANSS scores, as well as the positive and negative subscales of PANSS with overall Cowan's K. We calculated olanzapine equivalence scores for antipsychotic medication (Gardner, Murphy, O'Donnell, Centorrino, & Baldessarini, 2010). We performed Spearman bivariate correlations (2-tailed) in PSZ to examine the relationship between antipsychotic medication dose and overall Cowan's K in target trials (Table S3).

*Investigation of observed within group effects with age, IQ, psychopathology and medication:*

In order to investigate the possible influence of age, premorbid IQ, psychopathology and medication on the observed within group effects, we calculated several separate Spearman bivariate correlations (2-tailed) in each group. The effects were calculated as the difference in Cowan's K for flickering-bias/non-predictive cue and non-flickering-bias/non-predictive cue for the flickering effect, and non-flickering-bias/non-predictive cue and non-flickering-bias/predictive cue for the cue effect. The German Mehrfachwahl-Wortschatz-Intelligenz Test was administered to assess premorbid verbal intelligence (Lehrl, Merz, Burkhard, & Fischer, 2005), (Table S4).

*Catch trials*

In 20% of trials a non-target Gabor patch was probed at retrieval- for example when the instructions indicated that flickering information should be encoded ('flickering-bias', a non-flickering Gabor patch was probed. If a predictive cue was displayed, that cue indicated the incorrect locations. Overall accuracy across all four catch conditions was around chance level (PSZ = 55%, HCS = 57%, Table S1).

We conducted a LMM using catch trials to predict the WM score with group, salience, cue, age, and premorbid IQ (Formula: score ~ Salience * Cue * Group + Age + IQ). The model included subject as a random effect (formula: ~1 | Subject), (Table S5).

The post-hoc contrasts were computed using the Kenward-Roger degrees-of-freedom method with pairwise t-tests adjusted with the Tukey method and were adjusted for age and premorbid IQ. We investigated differences between (Table S6) and within groups (Table S7).

*Group differences for attentional prioritization*

Independent 2-tailed t-tests were conducted between groups for each condition to investigate possible differences in attentional prioritization (Cowan's K for target trials minus Cowan's K for catch trials, Table S8).

*Working memory capacity task:*

We implemented a 'canonical' color change detection task (Figure S1) on a personal computer using Presentation software in Version14.9 (www.neuro bs.com). Stimuli were presented on a grey background (RGB values: 191, 191, 191) in a dimly lit room with a viewing distance of approximately 60 cm. Throughout the experiment, a black fixation cross was displayed at the center of the screen. Each trial began with the alert phase, during which the fixation cross turned to red for 500 ms. This was followed by a preparation phase of 500 ms. During the encoding phase a sample array of four colored circles was presented for 200 ms. Each circle had a visual angle of approximately 0.95°. These circles were spaced equally apart on an imaginary circle with 12 possible locations around the black fixation cross covering a visual angle of approximately 5.25°, and the minimum distance between two circles was 0.29°. Each circle had one of seven easily discriminable possible colors with the following RGB values: black (0, 0, 0), red (255, 0, 0), white (255, 255, 255), blue (0, 0, 255), green (0, 255, 0), yellow (255, 255, 0), and magenta (255, 0, 255), with no repetitions of colors within a trial. During the delay phase, the black fixation cross remained on the screen for 1800 ms. A whole-display recognition test array followed, in which participants had a maximum duration of 3000 ms to decide if the test array was identical to the sample array presented in the encoding phase, or if one of the circles had changed color. Half of the trials were change trials (right mouse button), the other half no-change trials (left mouse button). In change trials, a randomly chosen circle changed its color. The total duration of each trial was 6000 ms followed by an inter-trial interval of 3000 ms. All participants received the same instructions prior to the beginning the task, and were asked to perform as accurately as possible, and to keep their eyes fixated constantly on the center of the screen. A total of 60 trials were tested in each participant, which required approximately nine minutes of testing time (Barnes-Scheufler et al., 2021).

**Figure S1. Working Memory Capacity Task**

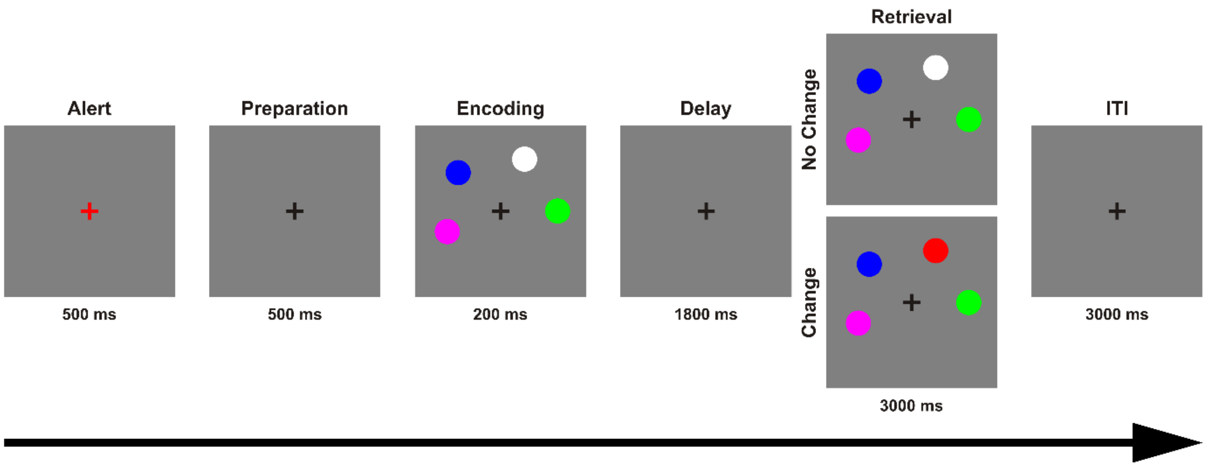

Figure S1. The change detection task used to assess working memory capacity. Each trial began with the alert phase, during which the fixation cross turned to red for 500 ms. This was followed by a preparation phase of 500 ms. During the encoding phase a sample array of four colored circles was presented for 200 ms. During the delay phase, the black fixation cross remained on the screen for 1800 ms. The whole-display recognition test array followed, in which participants had a maximum duration of 3000 ms to decide if the test array was identical to the sample array presented in the encoding phase, or if one of the circles had changed color. See (Barnes-Scheufler et al., 2021) for more details.

*Attentional prioritization and independent WM capacity estimate:*

We calculated Spearman correlations (two-tailed) of the independent WM capacity estimate (Pashler's K) and attentional prioritization efficiency (target Cowan's K – catch Cowan's K, Table S9).

*Possible effects of block:*

In order to detect performance changes across time, we correlated the sequential block number with block accuracy across both groups and within each group with Spearman correlations (two-tailed).

### RESULTS:

*Demographics:*

We compared handedness scores between groups using a t-test and did not find a significant difference ($t_{138} = 0.078$, $p = 0.938$).

We did not find a correlation between gender and overall target Cowan's K in PSZ ($r_s = 0.040$, $p = 0.752$), or in HCS ($r_s = 0.218$, $p = 0.062$).

#### Table S1. Accuracy

| | *Mean* | *SD* | *Range* |
|---|---|---|---|
| **HCS (*n* =74 )** | | | |
| *Global* | 0.77 | 0.07 | 0.55 – 0.90 |
| Flickering-bias/predictive cue | 0.78 | 0.08 | 0.55 – 0.93 |
| Flickering-bias/non-predictive cue | 0.78 | 0.08 | 0.52 – 0.93 |
| Non-flickering-bias/predictive cue | 0.77 | 0.07 | 0.56 – 0.89 |
| Non-flickering-bias/non-predictive cue | 0.76 | 0.08 | 0.49 – 0.89 |

| | | | |
|---|---|---|---|
| *Target* | 0.82 | 0.09 | 0.56 – 0.98 |
| Flickering-bias/predictive cue | 0.83 | 0.10 | 0.54 – 0.96 |
| Flickering-bias/non-predictive cue | 0.83 | 0.10 | 0.55 – 0.99 |
| Non-flickering-bias/predictive cue | 0.82 | 0.09 | 0.56 – 1.00 |
| Non-flickering-bias/non-predictive cue | 0.80 | 0.11 | 0.54 – 0.99 |
| *Catch* | 0.57 | 0.11 | 0.23 – 0.80 |
| Flickering-bias/predictive cue | 0.56 | 0.13 | 0.10 – 0.85 |
| Flickering-bias/non-predictive cue | 0.58 | 0.14 | 0.25 – 0.80 |
| Non-flickering-bias/predictive cue | 0.55 | 0.14 | 0.25 – 0.85 |
| Non-flickering-bias/non-predictive cue | 0.58 | 0.15 | 0.10 – 0.90 |
| **PSZ (*n* = 66)** | | | |
| *Global* | 0.70 | 0.08 | 0.54 – 0.84 |
| Flickering-bias/predictive cue | 0.72 | 0.07 | 0.54 – 0.87 |
| Flickering-bias/non-predictive cue | 0.70 | 0.10 | 0.48 – 0.88 |
| Non-flickering-bias/predictive cue | 0.70 | 0.09 | 0.49 – 0.91 |
| Non-flickering-bias/non-predictive cue | 0.66 | 0.09 | 0.48 – 0.83 |
| *Target* | 0.74 | 0.10 | 0.57 – 0.93 |
| Flickering-bias/predictive cue | 0.77 | 0.09 | 0.57 – 0.94 |
| Flickering-bias/non-predictive cue | 0.74 | 0.12 | 0.47 – 0.96 |
| Non-flickering-bias/predictive cue | 0.75 | 0.11 | 0.47 – 0.99 |
| Non-flickering-bias/non-predictive cue | 0.69 | 0.11 | 0.47 – 0.89 |
| *Catch* | 0.55 | 0.10 | 0.34 – 0.81 |
| Flickering-bias/predictive cue | 0.53 | 0.11 | 0.25 – 0.75 |
| Flickering-bias/non-predictive cue | 0.55 | 0.12 | 0.30 – 0.85 |
| Non-flickering-bias/predictive cue | 0.53 | 0.15 | 0.20 – 0.90 |
| Non-flickering-bias/non-predictive cue | 0.56 | 0.13 | 0.30 – 0.90 |

Table S1. Results of accuracy of main working memory task reported in mean, standard deviation (SD) and range (minimum - maximum).

### Table S2. Cowan's K

| | *Mean* | *SD* |
|---|---|---|
| **HCS (*n* =74 )** | | |
| *Global* | | |
| Flickering-bias/predictive cue | 1.104 | 0.332 |
| Flickering-bias/non-predictive cue | 1.102 | 0.330 |
| Non-flickering-bias/predictive cue | 1.068 | 0.293 |
| Non-flickering-bias/non-predictive cue | 1.025 | 0.333 |
| *Target* | | |
| Flickering-bias/predictive cue | 1.318 | 0.386 |
| Flickering-bias/non-predictive cue | 1.294 | 0.407 |
| Non-flickering-bias/predictive cue | 1.283 | 0.370 |
| Non-flickering-bias/non-predictive cue | 1.203 | 0.429 |
| *Catch* | | |
| Flickering-bias/predictive cue | 0.246 | 0.535 |
| Flickering-bias/non-predictive cue | 0.332 | 0.555 |
| Non-flickering-bias/predictive cue | 0.208 | 0.569 |

| | | |
|---|---|---|
| Non-flickering-bias/non-predictive cue | 0.311 | 0.611 |
| **PSZ (*n* = 66)** | | |
| *Global* | | |
| Flickering-bias/predictive cue | 0.880 | 0.300 |
| Flickering-bias/non-predictive cue | 0.819 | 0.380 |
| Non-flickering-bias/predictive cue | 0.816 | 0.353 |
| Non-flickering-bias/non-predictive cue | 0.666 | 0.370 |
| *Target* | | |
| Flickering-bias/predictive cue | 1.066 | 0.356 |
| Flickering-bias/non-predictive cue | 0.970 | 0.463 |
| Non-flickering-bias/predictive cue | 0.990 | 0.435 |
| Non-flickering-bias/non-predictive cue | 0.770 | 0.443 |
| *Catch* | | |
| Flickering-bias/predictive cue | 0.130 | 0.459 |
| Flickering-bias/non-predictive cue | 0.218 | 0.482 |
| Non-flickering-bias/predictive cue | 0.121 | 0.616 |
| Non-flickering-bias/non-predictive cue | 0.248 | 0.506 |

Table S2. Cowan's K reported as global score scores (mixed target and catch), as well as separated in target and catch trials per conditions.

### Table S3. Influences of psychopathology and medication

| | $r_s$ | *p Value* |
|---|---|---|
| **PSZ (*n* = 66)** | | |
| PANSS Positive | 0.097 | 0.446 |
| PANSS Negative | -0.173 | 0.172 |
| PANSS General | -0.012 | 0.926 |
| PANSS Total | -0.063 | 0.619 |
| Olanzapine Equivalence Score | -0.193 | 0.121 |

Table S3: Results of correlations in people with schizophrenia = PSZ of overall Cowan's K. We did not observe any significant correlation between any of the PANSS measures or the olanzapine equivalence score with overall Cowan's K in PSZ.

### Table S4. Flickering and cue effect correlations

| | $r_s$ | *p Value* |
|---|---|---|
| **HCS (*n* =74 )** | | |
| *Flickering Effect* | | |
| Age | 0.071 | 0.546 |
| Premorbid IQ | 0.034 | 0.771 |
| *Cue Effect* | | |
| Age | 0.240 | 0.040* |
| Premorbid IQ | -0.035 | 0.766 |

| | | |
|---|---|---|
| **PSZ (*n* = 66)** | | |
| *Flickering Effect* | | |
| Age | -0.188 | 0.130 |
| Premorbid IQ | -0.254 | 0.039* |
| PANSS Pos | 0.061 | 0.630 |
| PANSS Neg | -0.112 | 0.378 |
| PANSS Gen | 0.014 | 0.912 |
| PANSS Tot | -0.039 | 0.760 |
| Olanzapine Equivalence Score | -0.011 | 0.928 |
| *Cue Effect* | | |
| Age | -0.176 | 0.159 |
| Premorbid IQ | -0.127 | 0.310 |
| PANSS Pos | -0.048 | 0.709 |
| PANSS Neg | -0.037 | 0.772 |
| PANSS Gen | 0.022 | 0.861 |
| PANSS Tot | -0.036 | 0.779 |
| Olanzapine Equivalence Score | 0.018 | 0.887 |

Table S4: Results of correlations between demographic / clinical measures and the calculated flickering and cue effects. Interestingly, we observed a single significant correlation in HCS between the effect of cue and age and in PSZ between the effect of salience and premorbid IQ. Asterisks indicate significance $p < 0.001$ = ***, $p < 0.01$ = **, $p < 0.05$ = *.

**Figure S2. Results of catch trials**

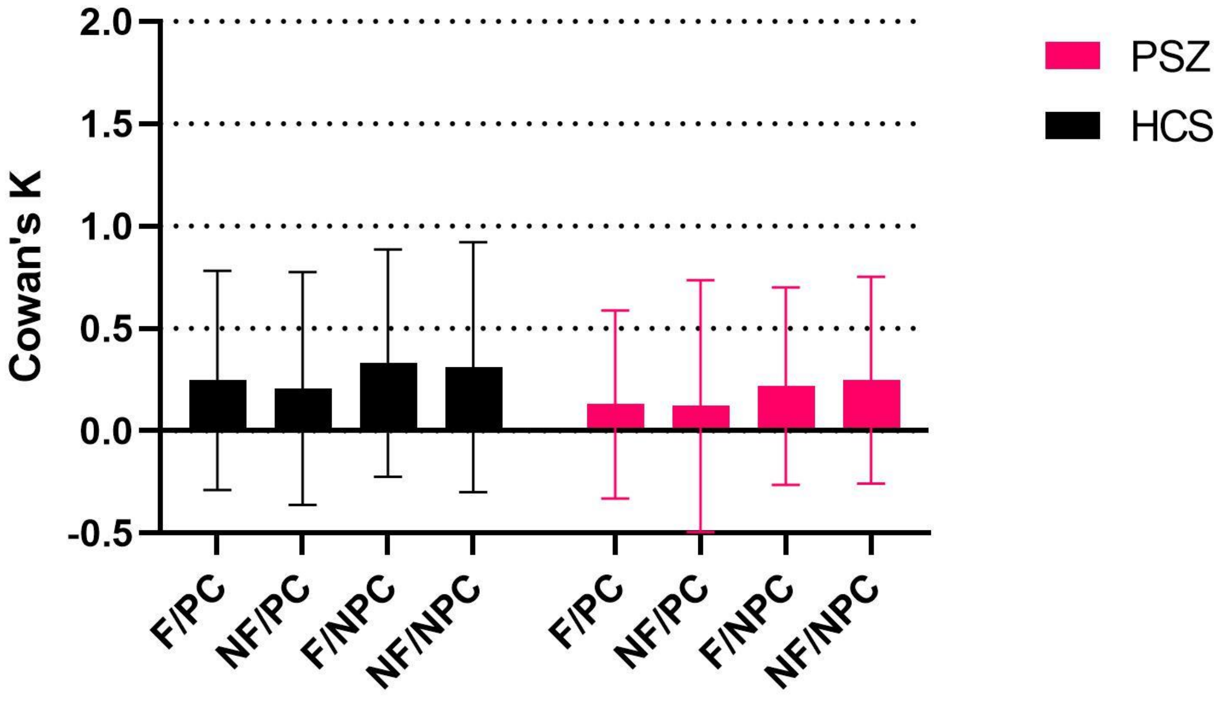

Figure S2. Amount of information stored in VWM in catch trials, estimated with Cowan's K in healthy control subjects = HCS and people with schizophrenia = PSZ. F/PC = flickering-bias/predictive cue, NF/PC = non-flickering-bias/predictive cue, F/NPC = flickering-bias/non-predictive cue, NF/NPC = non-flickering-bias/non-predictive cue. Error bars indicate standard deviation. We observed solely a significant effect of cue in the catch trials ($p = 0.004$). There were no significant effects of group, salience, age, or premorbid IQ (Table S5). Furthermore, there were no significant interactions of group by salience, group by cue, salience by cue, and no three-way interaction of group by salience by cue.

### Table S5. Results of LMM with catch trials

| | *df* | *F* | *p Value* |
|---|---|---|---|
| Group | 140 | 1.97 | 0.162 |
| Salience | 420 | 0.08 | 0.783 |
| Cue | 420 | 8.50 | 0.004** |
| Age | 140 | 0.01 | 0.925 |
| Premorbid IQ | 140 | 0.11 | 0.738 |
| Group x Salience | 420 | 0.34 | 0.561 |
| Group x Cue | 420 | 0.04 | 0.852 |
| Salience x Cue | 420 | 0.16 | 0.690 |
| Group x Salience x Cue | 420 | 0.03 | 0.867 |

Table S5: Results of LMM of catch trials with factors salience and cue, and covariates age and premorbid IQ. We observed solely a significant effect of cue in the catch trials. Asterisks indicate significance $p < 0.001$ = ***, $p < 0.01$ = **, $p < 0.05$ = *.

### Table S6. Results of between group comparisons of catch trials

| | *t* | *df* | *p Value* |
|---|---|---|---|
| Flickering-bias/predictive cue | 1.280 | 365 | 0.201 |
| Flickering-bias/non-predictive cue | 1.265 | 365 | 0.210 |
| Non-flickering-bias/predictive cue | 0.973 | 365 | 0.331 |
| Non-flickering-bias/non-predictive cue | 0.710 | 365 | 0.500 |

Table S6: Results of post-hoc t-tests between groups in each condition in catch trials. We did not observe any significant difference between groups in the catch trial conditions.

### Table S7. Results of within group comparisons of catch trials

| | *t* | *df* | *p Value* |
|---|---|---|---|
| **HCS (*n* =74 )** | | | |
| Flickering-bias/predictive cue vs Flickering/non-predictive cue | -1.275 | 426 | 0.579 |
| Flickering-bias/predictive cue vs Non-flickering/predictive cue | 0.558 | 426 | 0.944 |
| Non-flickering-bias/predictive cue vs Non-flickering/non-predictive cue | -1.514 | 426 | 0.430 |
| Non-flickering-bias/non-predictive cue vs. Flickering/non-predictive cue | -0.319 | 426 | 0.990 |
| **PSZ (*n* = 66)** | | | |
| Flickering-bias/predictive cue vs Flickering/non-predictive cue | -1.224 | 426 | 0.612 |
| Flickering-bias/predictive cue vs Non-flickering/predictive cue | 0.127 | 426 | 0.999 |

| | | | |
|---|---|---|---|
| Non-flickering-bias/predictive cue vs Non-flickering/non-predictive cue | -1.772 | 426 | 0.288 |
| Non-flickering-bias/non-predictive cue vs. Flickering/non-predictive cue | 0.422 | 426 | 0.975 |

Table S7: Results of post-hoc t-tests within groups between conditions in catch trials. We did not observe any significant differences between conditions within groups.

### Table S8. Group differences for attentional prioritization

| | *t* | *df* | *p Value* |
|---|---|---|---|
| Flickering-bias/predictive cue | -1.312 | 138 | 0.192 |
| Flickering-bias/non-predictive cue | -1.756 | 138 | 0.081 |
| Non-flickering-bias/predictive cue | -1.580 | 138 | 0.116 |
| Non-flickering-bias/non-predictive cue | -2.880 | 138 | 0.005** |

Table S8: Results of post-hoc t-tests between groups in each condition in attentional prioritization (Cowan's K for target trials minus Cowan's K for catch trials). We observed a single significant difference between groups in the efficiency of attentional prioritization in the non-flickering-bias/non-predictive cue condition. Asterisks indicate significance $p < 0.001$ = ***, $p < 0.01$ = **, $p < 0.05$ = *.

### Table S9. Results of attentional prioritization and independent WM capacity estimate

| | $r_s$ | *p Value* |
|---|---|---|
| **HCS (*n* =74 )** | | |
| Flickering-bias/predictive cue | .189 | 0.108 |
| Flickering-bias/non-predictive cue | .155 | 0.187 |
| Non-flickering-bias/predictive cue | .140 | 0.234 |
| Non-flickering-bias/non-predictive cue | .148 | 0.209 |
| **PSZ (*n* = 66)** | | |
| Flickering-bias/predictive cue | .090 | 0.473 |
| Flickering-bias/non-predictive cue | .101 | 0.421 |
| Non-flickering-bias/predictive cue | .101 | 0.421 |
| Non-flickering-bias/non-predictive cue | .212 | 0.087 |

Table S9. Results of Spearman correlations of independent WM capacity estimate (Pashler's K) and attentional prioritization efficiency (target Cowan's K – catch Cowan's K). We did not find any significant correlations.

#### *Possible Effects of Block:*

We did not find any effects of block across groups ($r_s$ = -0.006, $p$ = 0.776) as well as for PSZ ($r_s$ = 0.029, $p$ = 0.314), or HCS ($r_s$ = -0.050, $p$ = 0.096). Thus, performance did not improve or degrade over time.
